# *Coxiella burnetii* epitope-specific T-cell responses in chronic Q fever patients

**DOI:** 10.1101/582007

**Authors:** Anja Scholzen, Guilhem Richard, Leonard Moise, Eva Hartman, Chantal P. Bleeker-Rovers, Patrick M. Reeves, Susan Raju Paul, William D. Martin, Anne S. De Groot, Mark C. Poznansky, Ann E. Sluder, Anja Garritsen

## Abstract

Infection with *Coxiella burnetii*, the causative agent of Q fever, can result in life-threatening persistent infection. Reactogenicity hinders worldwide implementation of the only licensed human Q fever vaccine. We previously demonstrated long-lived immunoreactivity in individuals with past symptomatic and asymptomatic *Coxiella* infection (convalescents) to promiscuous HLA-class II *C. burnetii* epitopes, providing the basis for a novel T-cell-targeted subunit vaccine. Here we investigated in a cohort of 22 individuals with persistent infection (chronic Q fever) whether they recognize the same set of epitopes, or distinct epitopes that could be candidates for a therapeutic vaccine or aid in the diagnosis of persistent infection.

Individuals with chronic Q fever showed strong class II epitope-specific cultured ELISpot responses largely overlapping with the peptide repertoire identified previously for convalescents. Five additional peptides were recognized more frequently by chronic subjects, but there was no combination of epitopes uniquely recognized by or non-reactive in chronic Q fever subjects. Consistent with more recent/prolonged exposure, we found, however, stronger direct *ex vivo* responses to whole-cell *C. burnetii* and individual peptides in direct ELISpot than in convalescents.

In conclusion, we have validated and expanded a previously published set candidate epitopes for a novel T-cell targeted subunit Q fever vaccine in the context of chronic Q fever patients and demonstrated that they successfully mounted a T-cell response comparable to that of convalescents. Finally, we demonstrate that individuals treated for chronic Q fever mount a broader *ex vivo* response to class II epitopes than convalescents, which could be explored for diagnostic purposes.

## Introduction

Q fever is a zoonotic disease that is endemic in many countries worldwide. It is caused by the environmentally highly stable small Gram-negative coccobacillus *Coxiella burnetii*, which is transmitted to humans predominantly by aerosol from infected ruminants such as goats, sheep and cattle (1). Outbreaks usually occur in the occupational setting including the livestock industry and deployed military personnel (1). *Coxiella* outbreaks can also occur in the general population, the largest to date being the outbreak in the Netherlands from 2007-2010 with an estimated 40,000 infections at the center of the epidemic area alone (2). While infection remains asymptomatic in an estimated 50-60% of individuals and acute symptomatic infection is readily treatable with antibiotics, a large proportion (10-20%) of individual with acute Q fever later develop Q fever fatigue syndrome. Further, 1-5% of (often asymptomatically) infected individuals progress to persistent infection also known as chronic Q fever. Chronic Q fever has a poor prognosis and manifests as endocarditis, infected aneurysms or vascular prosthesis infection in individuals with specific risk factors (1, 3).

While Q fever infection in humans can be prevented by vaccination using Q-VAX®, an inactivated whole cell vaccine based on phase I *C. burnetii*, it is licensed for use in Australia only. Importantly, this vaccine requires pre-vaccination screening for prior exposure due to reported side effects in previously exposed individuals (3–5). In this context, the objective of the Q-VaxCelerate consortium is to develop a novel non-reactogenic T-cell-targeted human Q fever vaccine that does not require pre-screening of vaccinees, rationally selecting T-cell epitopes for inclusion in such a vaccine (5, 6). Using immunoinformatically predicted T-cell epitopes derived from *C. burnetti* sero-reactive and type IV secretion systems (TSS4) substrate proteins, we previously analyzed antigenicity in naturally infected subjects with past symptomatic or asymptomatic *C. burnetii* infection; these are here-after referred to as ‘convalescents’ since infection was cleared. In these naturally exposed subjects, we demonstrated long-lived immunoreactivity to promiscuous CD4 T-cell epitopes, while HLA class I epitope responses were sparse in this cohort (7). One possible explanation for the latter was that class I responses might have contracted faster than class II responses, as previously observed following smallpox infection or vaccination and tuberculosis treatment (8–11). In this initial study, there were no striking differences between past asymptomatic or symptomatic infected individuals, all of whom successfully cleared acute *C. burnetii* infection. The question remains, however, as to whether analogous to herpes simplex virus infection, there might be distinct epitope-specific T-cell repertoires for individuals that either successfully control infection or develop persistent infection (12). Such epitopes might be interesting targets for a potentially separate therapeutic vaccine to accelerate bacterial clearance in chronic Q fever, or aid in the diagnosis of this persistent infection.

In the present study we therefore analyzed T-cell reactivity to the same set of epitopes in a cohort of subjects diagnosed and treated for persistent *C. burnetii* infection (chronic Q fever). The aim was to investigate whether subjects with chronic Q fever (i) show potentially greater reactivity to class I epitopes given their more recent exposure, (ii) recognize the same or a distinct set of class II epitopes and (iii) differ in their effector memory T-cell response profile compared to individuals with resolved acute symptomatic or past asymptomatic infection.

## Results

### Chronic Q fever subjects have cultured ELISpot response patterns to HLA class I and II *C. burnetii* epitopes comparable to that of convalescent subjects

A group of 22 individuals with proven (n=16) and probable (n=6) chronic Q fever consented for participation in this study (Table 1). All but two chronic Q fever subjects still had phase I IgG titers of ≥1024 at inclusion into the study (median with interquartile range: 4096 [1536-8192]), and 13/16 proven and 1/6 probable subjects were still undergoing antibiotic treatment. For analysis of T-cell epitope-specific responses in the chronic Q fever cohort, preference was given to individuals with proven chronic Q fever who were diagnosed in 2016 or later and still undergoing antibiotic treatment. Subjects were scheduled for blood collection based on availability for class I and II peptide screening (Table 1). In total, 13 proven and one probable chronic Q fever patients were tested for promiscuous class II epitope-specific responses (Table S1), and 10 proven and three probable chronic Q fever patients for class I epitope-specific responses (Table S2). HLA typing of the two selected groups showed supertype distributions largely comparable to expected frequencies in the general population and/or those in the previously analyzed convalescent groups (Table S3 and S4), except for an underrepresentation of HLA-DR11, A11 and B8 and an overrepresentation of HLA-A3 supertypes, which may be partially attributed to the small group sizes.

**Table 1.**
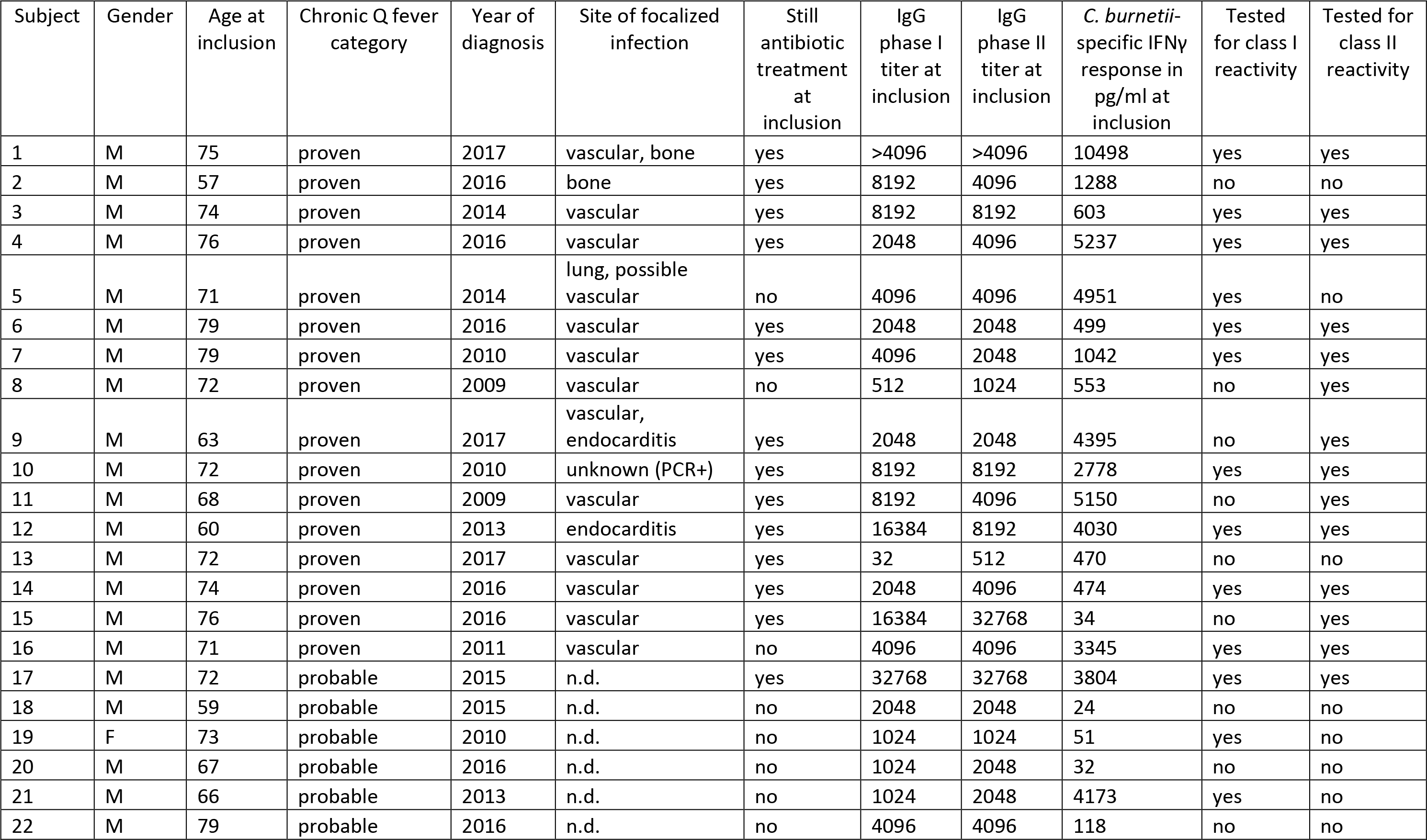
Chronic Q fever cohort

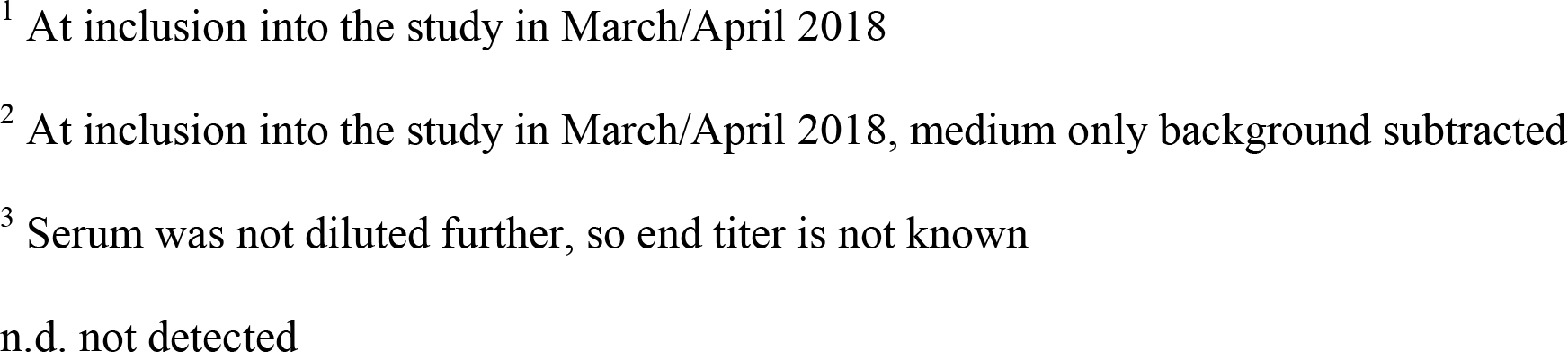

T-cell responses were first analyzed by cultured IFNγ ELISpot, which both enhances detection of low frequency responses and preferentially measures central memory T-cells (13). Similar to previous observations for convalescent *C. burnetii*-exposed subjects, 9/14 chronic Q fever patients (64%) showed responses to 3-14 HLA class II peptides per donor (Figure 1A), while responses to HLA class I peptides were rare, with only three subjects showing responses to one or two peptides each (Figure 1B). When directly comparing the data from the chronic Q fever cohort to convalescent subjects (asymptomatic, n=33; and symptomatic, n=23), there was no statistically significant difference in the breadth of the class II response per subject between either all chronic and all convalescent subjects (p=0.15 by Mann-Whitney test), or chronic subjects on the one hand and convalescent symptomatic or asymptomatic subjects on the other hand (p=0.90 and p=0.16 by Kruskal-Wallis test with Dunn’s multiple comparison post hoc test) (Figure 2A). Nevertheless, chronic subjects had the smallest proportion of non-responders amongst the three groups (Figure 2B).

**Figure 1.**
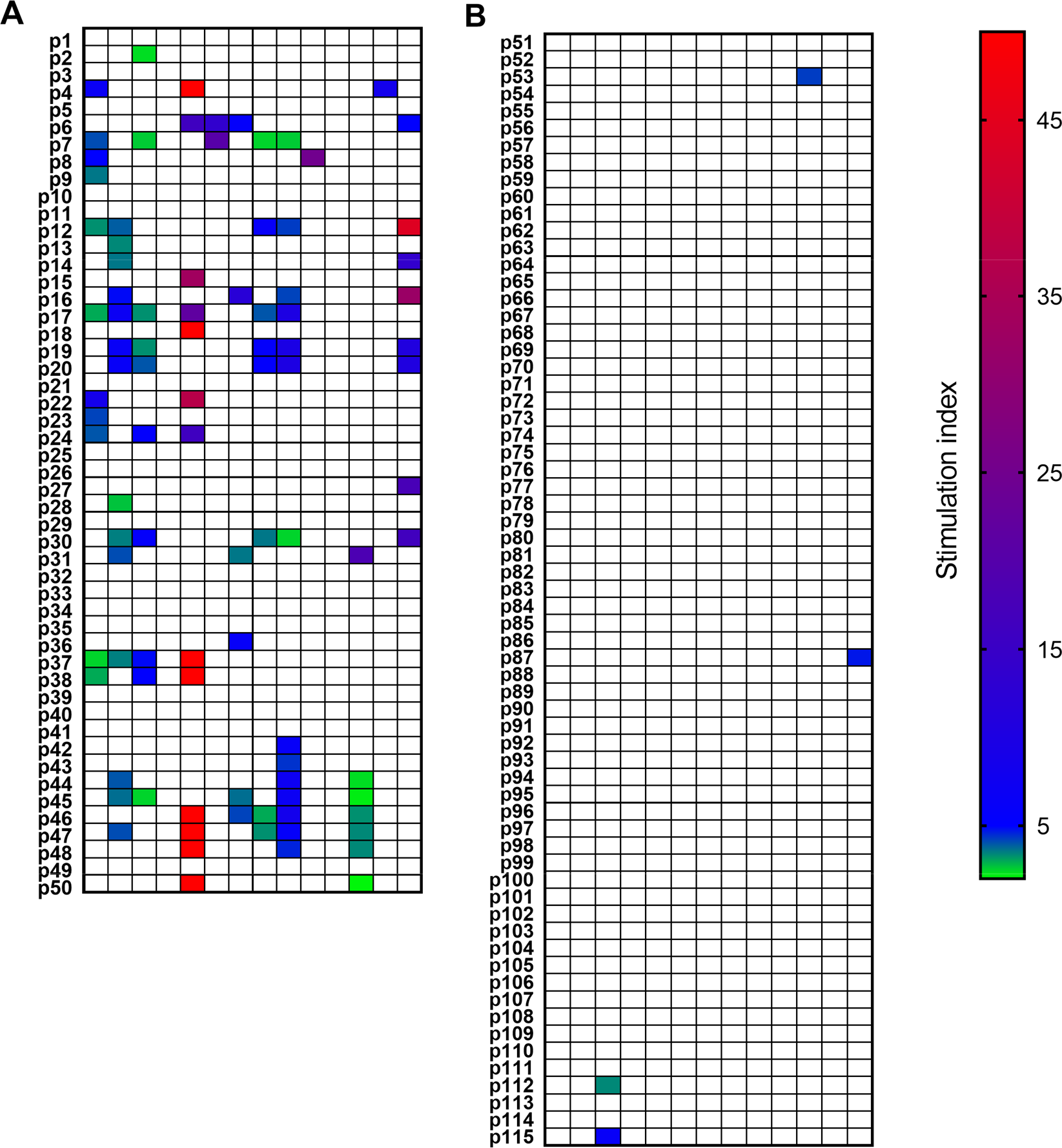
Cultured ELISpot human IFNγ responses to HLA class I and II peptides in individuals treated for chronic Q fever. Individual IFNγ responses to (A) HLA class II and (B) class I peptides determined by cultured ELISpot are depicted as stimulation indices (SI). Each column shows data from one donor, each row responses to one of the 50 class II or 65 class I peptides. Responses not significantly different from background and/or lower than an average of 10 spots/well are denoted as 0. Significant responses with a SI≥2 are color coded as per heatmap legend. Responses for one donor were capped at SI=50 to be able to properly resolve the magnitude of responses of the remaining subjects.

**Figure 2.**
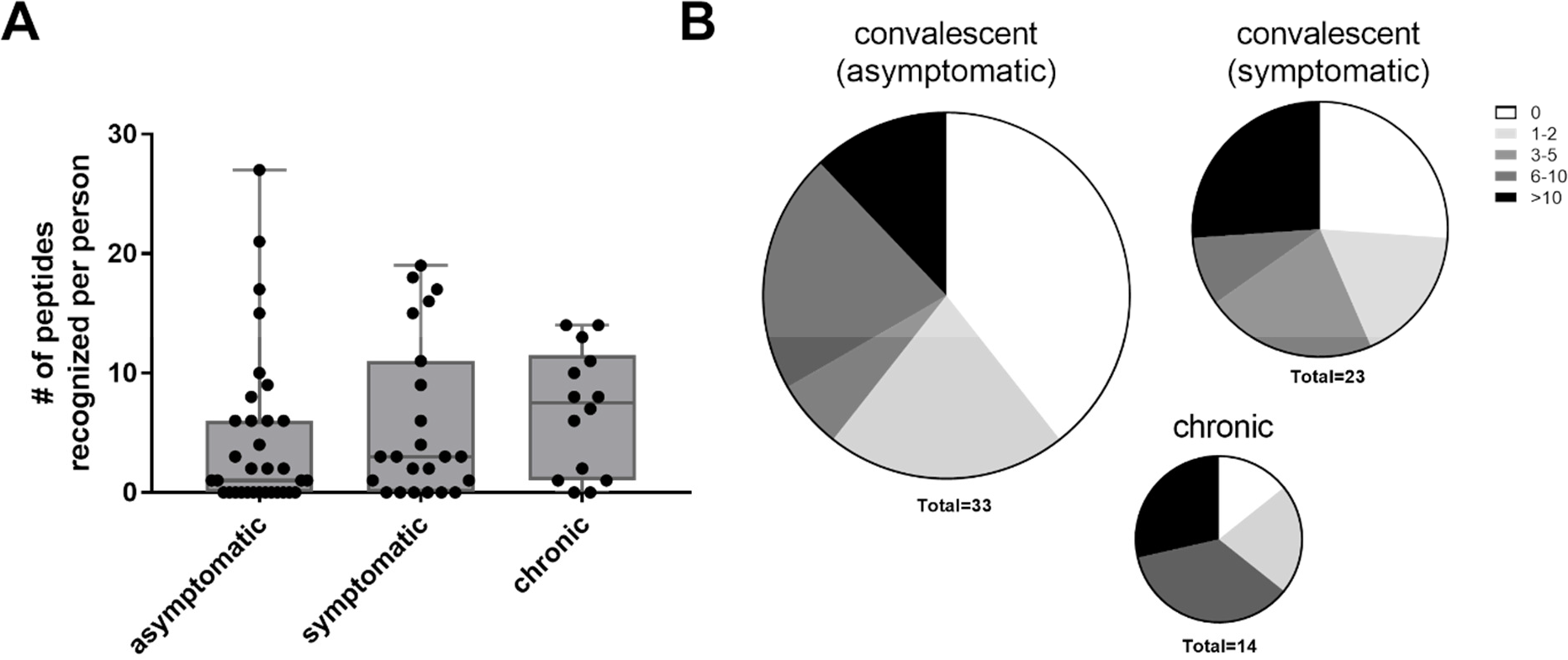
Cumulative HLA class II peptide responses in chronic compared to convalescent individuals. Data are shown for past asymptomatic (n=33) or symptomatic (n=23) infected individuals and for chronic Q fever subjects (n=14) as the cumulative peptide response (SI≥2) per donor (A) or as the proportion of subjects recognizing 0, 1-2, 3-5, 6-10 or >10 peptides (B). Whisker-dot-plots show the median and interquartile range (25^th^ and 75^th^ percentile) with whiskers extending from min to max values.

Both the overall breadth of responses and the responses to individual class II peptides largely overlapped between chronic and convalescent subjects: In total, 33/50 HLA class II peptides were recognized by at least 1/14 chronic subjects; comparable to the fraction (28/50) of HLA class II peptides recognized by a similar proportion (at least 4/56, 7.14%) of convalescent individuals. The same 22 peptides were recognized in both cohorts by at least 7% of the subjects. The peptides that were not recognized by any individual in the chronic cohort included 5/6 peptides that were also not recognized by any convalescent subject (p11, p34, p35, p40 and p49, Figure 3).

**Figure 3.**
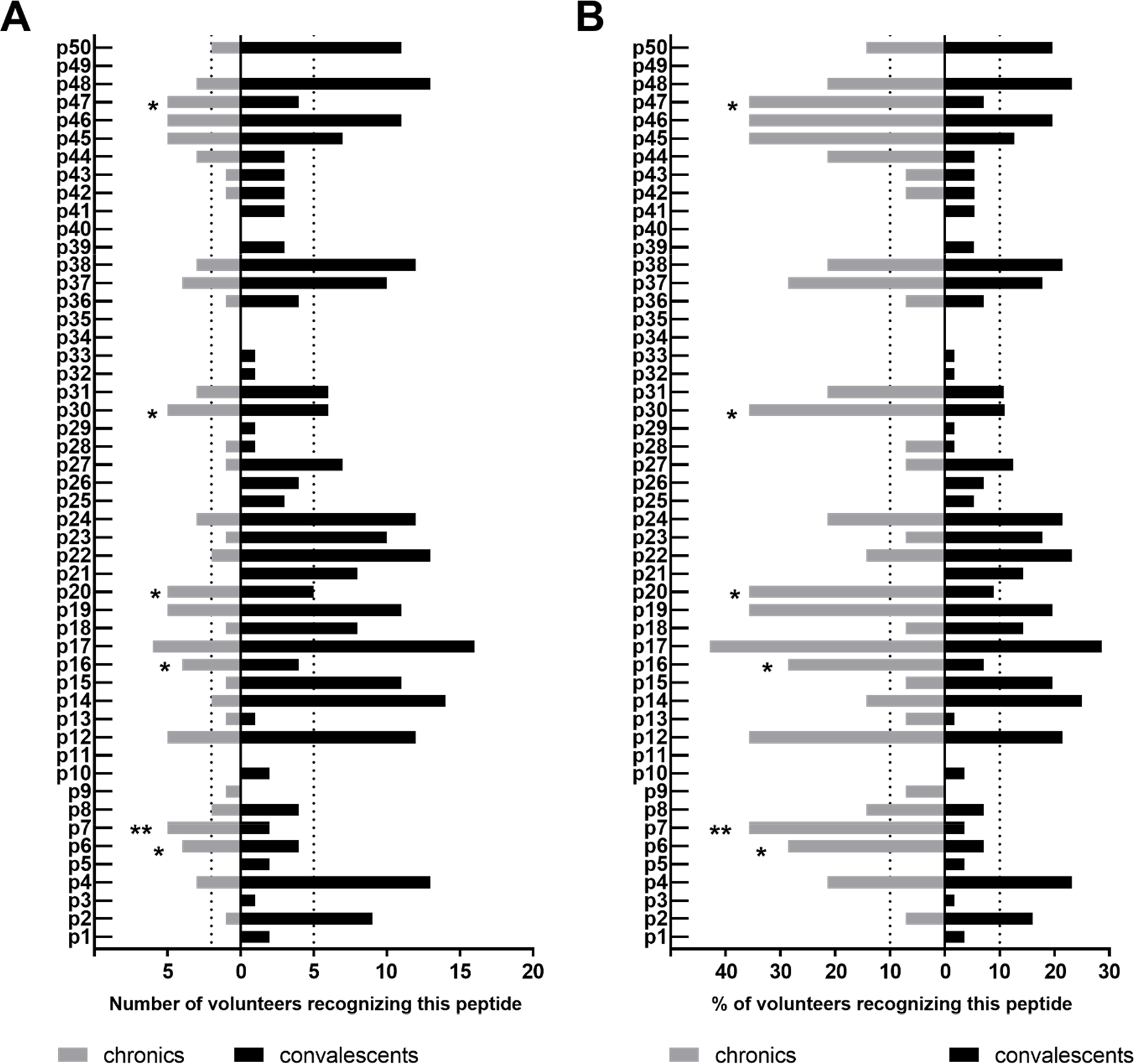
Class II peptide antigenicity patterns in chronic and convalescent individuals. Data are shown as the number (A) and proportion (B) of individuals with IFNγ responses to the 50 individual peptides in the chronic (n=14; grey bars) and convalescent cohorts (both past asymptomatic and symptomatic; n=56; black bars), as determined by cultured ELISpot. Bars extending over dotted lines indicate those peptides that were recognized by more than 10% of chronic (≥2/14) or convalescent subjects (>5/56). Asterices indicate significant difference in proportion between the two groups by Fisher’s exact test. * p<0.05; ** p<0.01

More importantly, out of 21 highly antigenic HLA class II peptides that were previously found to be recognized by >10% of all convalescent individuals (at least 6/56), 15 were also recognized by >10% of chronic subjects (at least 2/14 and up to 6/14 = 42%; Figure 3). This included at least one epitope from each of the five source proteins that were represented by two highly antigenic epitopes each in the convalescent cohort (p14+p15, CBU_1835/protoporphyrinogen oxidase; p18+p19, CBU_1513/protoporphyrinogen oxidase; p22+p23, CBU_1398/SucB, p37+p38, CBU_0718, p45+p46, CBU_0307/outer membrane protein). Another five of these 21 highly antigenic peptides were recognized by 1/14 chronic subjects. Only a single peptide that was highly antigenic in convalescents (p21 from CBU_1416/repressor protein C2), was not recognized by any chronic subject tested; however, 5/14 chronic individuals did recognize a second peptide (p2) from the same source protein.

Many of the class II peptide responses were at least as frequent in chronically infected subjects as in convalescents, and despite the large difference in group sizes, responses to six class II peptides were statistically significantly more frequent in chronic subjects (p6, p7, p16, p20, p30 and p47; Figure 3). All of these six peptides were recognized by four to five individuals (28-36%) within the chronic cohort, while for five of these peptides (all but p30), the frequency of responses in the convalescent cohort was <10% (two to five out of 56 subjects). However, for 3/5 peptides to which >10% chronic subjects but <10% convalescents reacted, >10% convalescents did show a response to a second epitope from the same source protein. Finally, there was only a single peptide (p9, the single screened epitope from the hypothetical exported protein CBU_2065) that was recognized by one chronic subject but not a single individual in the convalescent cohort. Taken together, while (central memory) cultured ELISpot responses to some individual peptides are more frequent in chronic subjects, there is no strong evidence for a set of source proteins or epitopes for which responses are uniquely present or absent in subjects with chronic Q fever.

### Chronic Q fever subjects show more frequent direct ELISpot responses to HLA class II *C. burnetii* epitopes than convalescent subjects

Given that subjects treated for chronic Q fever had a more recent and prolonged exposure to *C. burnetii*, we hypothesized that these individuals might also show a stronger effector memory T-cell response profile compared to individuals with resolved acute Q fever or past asymptomatic infection. Indeed, the individuals with chronic Q fever enrolled in this study showed significantly higher IFNγ secretion measured following whole blood stimulation with heat-killed whole-cell *C. burnetii* (strain Cb02629) compared to convalescent subjects (Table 1, Figure S1A). Stimulation of freshly isolated PBMCs for direct ELISpot with whole-cell *C. burnetii* indicated that this was at least partially due to a higher frequency of responding cells, with significantly higher numbers of spot forming units and higher stimulation indices in chronic compared to convalescent subjects (Figure S1B-C). Multiplex cytokine analysis of supernatants from whole blood stimulation revealed that the greater *ex vivo* response was not just confined to IFNγ, but also evident for IL-2 responses to whole-cell heat-killed *C. burnetii* (Figure S2A). The ratio between IFNγ and IL-2 responses in chronic subjects did not differ from that found for convalescent subjects (Figure S2A) IL-10 responses, in contrast, were lower in the chronic Q fever cohort, and innate TNFα and IL-1β responses did not differ between chronic and convalescent subjects (Figure S2B).

For a subset of chronic and convalescent individuals we next analyzed by direct ELISpot whether HLA class II *C. burnetii*-specific peptide responses would also be more readily detected in chronic patients. This assay preferentially measures effector memory responses (13). The proportion of individuals with direct ELISpot responses in both cohorts was comparable, with 5/11 responding convalescent individuals and 7/13 chronic subjects (Figure 4). However, the breadth of the response was larger for chronic subjects: one of the seven responding chronic subject showed *ex vivo* responses to four class II peptides and another four subjects scored positive for 6-12 class II peptides. In contrast, only one of the responding convalescent subjects recognized four peptides and the remaining three individuals only one to two peptides. All but one chronic subject and all convalescent individuals with detectable responses by direct ELISpot also showed responses by cultured ELISpot, and the individual peptides recognized in both assays per donor largely overlapped between these two groups (Figure S3 and S4). Although there were three peptides that elicited direct re-call responses with a relatively high proportion of individuals exclusively in the chronic group (p7 in 4/13; p10 and p19 in 3/13), this difference did not reach statistical significance by Fisher’s exact test, given the small number (n=11) of convalescent subjects also tested. Amongst convalescent subjects, the only peptides recognized by more than one individual were p4 and p38, two of the highly reactive peptides in the cultured ELISpot assay.

**Figure 4.**
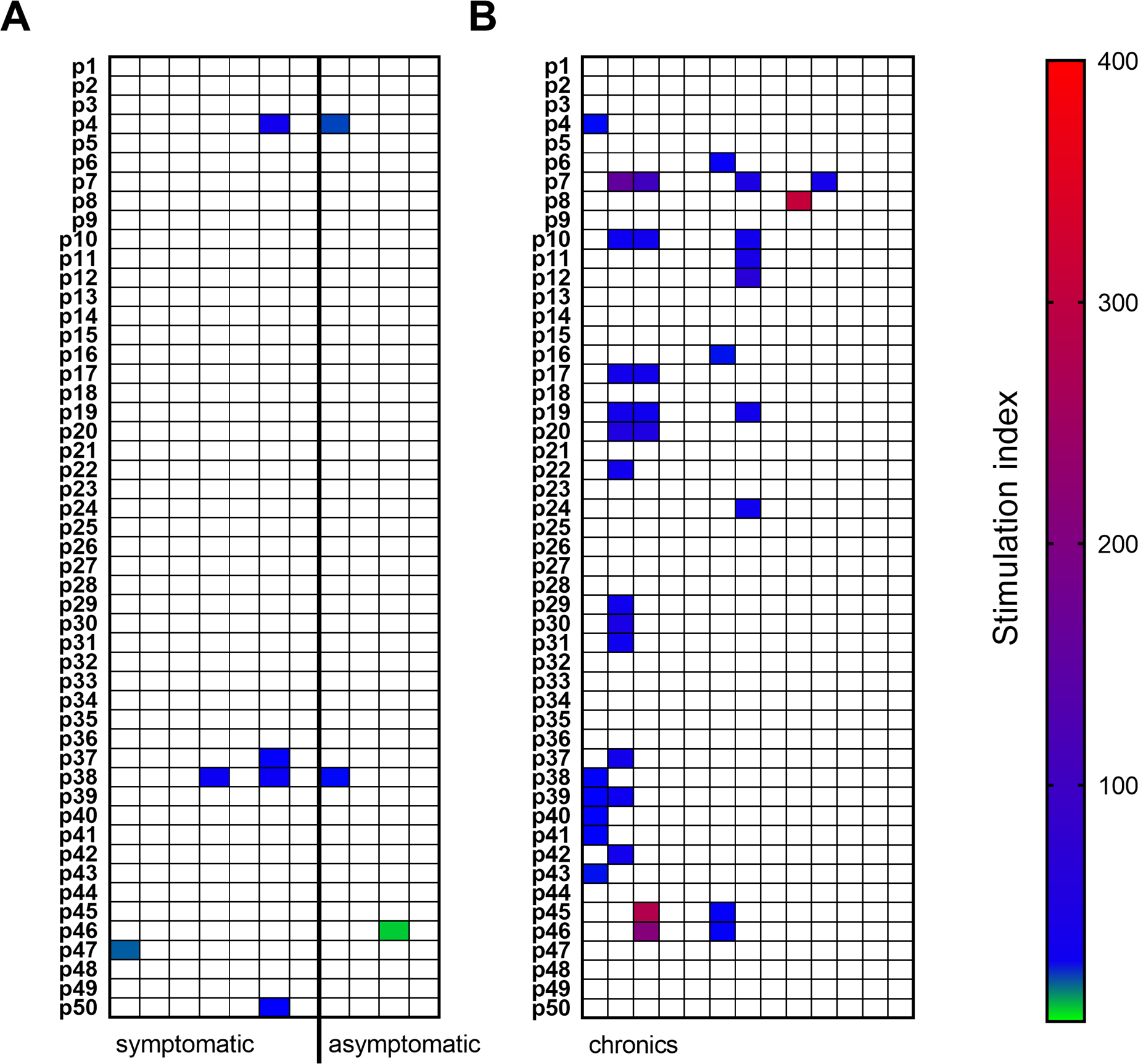
Direct ELISpot human IFNγ responses to HLA class II peptides. IFNγ responses to individual HLA class II peptides were determined by direct ELISpot for (A) convalescent individuals with a past history of symptomatic (n=7) or asymptomatic (n=4) Q fever infection and (B) individuals with chronic Q fever (n=13). Data are depicted as stimulation indices (SI) for all subjects analyzed. Each column shows data from one donor, each row responses to one of the 50 class II peptides. Responses not significantly different from background and/or lower than an average of 10 spots/million cells plated are denoted as blanks. Significant responses with a SI≥2 are color coded as per heatmap legend.

## Discussion

In this study we compared the repertoire of HLA class II T-cell epitopes recognized by subjects treated for persistent *C. burnetii* infection (chronic Q fever) with those recognized by convalescent individuals (i.e. those with resolved past acute or asymptomatic infection) in our previous study (7). We find that individuals treated for chronic Q fever have effectively generated a central memory *C. burnetii*-specific T-cell response, as measured by cultured ELISpot, that closely resembles that of convalescent patients. This includes both the breadth and the individual HLA class II epitopes recognized, as well as the near absence of detectable responses to HLA class I peptides. The main differences between the two cohorts were that compared to convalescents chronic Q fever subjects showed a higher proportion of cultured ELISpot responses to a small subset of six promiscuous CD4 T-cell epitopes, and exhibited detectable effector memory responses, as measured by direct ELISpot, to a greater number of peptides per subject. Both are consistent with more recent and prolonged antigen exposure in the chronic Q fever subjects.

Our study provides validation of the highly antigenic potential of the previously identified 22 promiscuous HLA class II peptides and the identification of an additional five *C. burnetii* HLA class II epitopes (p6, p7, p16, p20 and p47). These five peptides were recognized by four to five individuals (28-36%) within the chronic cohort, highlighting their strong antigenic potential at least during/ shortly after infection. Bearing in mind the small size of the group of chronic patients evaluated, the fact that all but one of the epitopes found to be highly antigenic in convalescent subjects (recognized by >10% individuals) were also recognized by at least one subject in the chronic cohort further indicates that at least amongst this set of screened HLA class II epitopes, there is no unique set of peptides to which responses would be absent in persistently infected individuals, and that would warrant consideration for a separate therapeutic vaccine for chronic Q fever patients. These results for chronic Q fever are in contrast to the observation of “asymptomatic” epitopes in herpes simplex virus infection (12). The principle difference may be that unlike Q fever, herpes simplex infection is always considered a persistent infection but can remain asymptomatic nonetheless. Instead, the same set of promiscuous HLA class II epitopes identified previously in the convalescent cohort (7) could in principle be used to further boost already primed T-cell responses in individuals with persistent infection. Whether this would speed up resolving of infection in this patient group, however, is unclear. Evidently, the IFNγ re-call response of circulating T-cells in individuals with chronic Q fever is fully functional – both in response to individual epitopes and whole-cell *C. burnetii*. This is in line with previous studies using IFNγ ELISA following whole blood stimulation (14, 15) and ELISpot using freshly isolated PBMCs (16). If this strong IFNγ response is insufficient to promote clearance of infection foci by activating *C. burnetii*-infected monocytes/macrophages then the defect could be downstream of IFNγ signaling as proposed previously (14). In particular, antigen-presenting cell maturation, function and interaction with T-cells as mediated via the IFN-IL-12p40 feedback loop (14, 17–19) in foci of infection could be compromised in persistent infection. Therefore, further research is required to clarify whether chronic Q fever patients can benefit from a therapeutic vaccine, or whether a completely different approach is needed to achieve clearance in this population.

The results of this study indicate that the unexpected scarcity of detectable responses to the predicted HLA class I epitopes in convalescent subjects (7) is not simply due to the long time-lapse between initial exposure and T-cell assays in our previous study, given that the chronic subjects analyzed here were exposed to *C. burnetii* antigens until much more recently. An obvious potential confounder in this chronic Q fever cohort is the fact that these subjects had been diagnosed and received antibiotic treatment for various lengths of time. Heterogeneity in time since diagnosis and in the duration of ongoing treatment, however, was minimized during selection of individuals for epitope screening. Moreover, a previous study showed that duration of antibiotic treatment following diagnosis of chronic Q fever, and whether subjects received treatment or not, did not influence IFNγ secretion, at least not in a whole blood stimulation assay using whole-cell *C. burnetii* (14). Of note, CD8 responses have been shown to decline rapidly following *Mycobacterium tuberculosis* treatment (10, 11) and we cannot exclude that this might have also impacted class I responses in the present cohort. The question whether and which class I epitopes should be included in a T-cell targeted Q fever vaccine for humans therefore requires further investigation during a new outbreak or a vaccination campaign.

In line with the observed higher frequency of circulating effector T-cell responding directly *ex vivo* to *C. burnetii* epitopes as well as whole-cell *C. burnetii* in these persistently infected individuals, we not only find stronger IFNγ responses, but also significantly higher IL-2 production in chronic compared to convalescent individuals. IL-2 is mainly produced by antigen-specific activated CD4 T-cells shortly after T-cell receptor engagement (20). Of note, these results contrast with a previous study that found lower IL-2 secretion in response to whole-cell *C. burnetii* and elevated IFNγ/IL-2 ratio in patients with chronic Q fever. This finding was hypothesized to reflect increased numbers of circulating effector T-cells producing IFNγ and low amounts of IL-2 (14). However, a simple supernatant secretion assay does not distinguish whether different cytokines are produced by the same or different cell populations, and central memory T-cells can also co-produce two or more cytokines including IL-2 (21). A possible technical explanation for the discrepancy between the two studies in regards to IL-2 secretion and IFNγ/IL-2 ratio is that in the previous study IL-2 responses were assessed after 48 h rather than 24 h stimulation (14), which may have impacted measurement of this rapidly consumed growth factor. Moreover, while individuals in both studies were recruited from the same region and all were likely initially exposed during the 2007-2011 outbreak, patients for the other study were assessed approximately 5 years earlier. Thus, cellular responses assessed herein have likely further contracted in convalescents since their initial exposure, potentially to different degrees for each cytokine. Infection and thus antigen exposure in chronic patients, in contrast, was persistent and hence longer. Whether the longer clinical pre-patency in our cohort might potentially relate to stronger IL-2 responses can only be speculated and requires investigation in a larger, specifically designed study.

One obvious question arising from this limited dataset is whether direct *ex vivo* responses to specific epitopes such as p7, p10, and p19, which were confined to the chronic cohort, might be of diagnostic value. This would require evaluation of a much larger cohort of past exposed individuals and (ideally recently diagnosed) chronic Q fever cases and a separate group of subjects with acute Q fever or recently recovered individuals. Only then will it be possible to determine whether *ex vivo* responses to these epitopes correlate with persistent infection, or simply with recent exposure. To be of value as a diagnostic tool, coverage of subjects would have to be greater than the currently observed 23-30%. Otherwise, the assay would at best be of supporting value in addition to the existing set of PCR, serology and scanning techniques to localize infection.

In conclusion, we herein validate and expand the characterization of a previously published set of promiscuous *C. burnetii*-specific HLA class II T-cell epitope clusters as candidates for a novel T-cell targeted subunit Q fever vaccine. We find that chronic Q fever patients have mounted a central memory re-call T-cell response comparable to that of convalescent individuals. Finally, we demonstrate that individuals treated for chronic Q fever mount a broader *ex vivo* response to class II epitopes, which could be explored for diagnostic purposes.

## Materials and Methods

### Study population

Twenty-two participants were recruited who were receiving or had received treatment for chronic Q fever or persistent focalized infection at the outpatient clinics of the Radboud university medical center in Nijmegen, the Elisabeth Hospital in Tilburg, the Jeroen Bosch Hospital in ’s-Hertogenbosch, the Bernhoven Hospital in Uden, the Medisch Spectrum Twente hospital in Twente, the Medical University Centre in Maastricht and the Zuyderland Medical Center in Heerlen, the Netherlands. The chronic Q fever group comprised 16 proven and 6 probable chronic Q fever patients diagnosed according to the Dutch consensus guideline on chronic Q fever (22). Ten of the proven chronic Q fever patients presented with a vascular focus in an aneurysm or aortic prosthesis, one with endocarditis, one with both a vascular and valvular localization, one with both vascular and vertebrae foci, one with lesions in the lung and an aneurysm, one with a focus in vertebrae only and one with a positive PCR. Thirteen of the proven patients and one of the probable cases were still on antibiotic treatment at the time of inclusion into this study (Table 1). At inclusion into the study, all participants treated for chronic Q fever completed a medical questionnaire and donated blood for HLA typing and analysis of serological and cellular responses to whole-cell *C. burnetii*.

*C. burnetii*-specific cultured ELISpot responses determined for this chronic Q fever cohort were compared to previously published results (7) from past exposed individuals with a history of resolved asymptomatic or symptomatic Q fever infection who recruited from the village of Herpen, the Netherlands, one of the focal centers of the 2007-2010 Q fever outbreak (23, 24). Additional assays presented in the present study (direct ELISpot responses and cytokine release during whole blood stimulation) were conducted using a subgroup from this cohort of convalescent Q fever exposed individuals from Herpen.

The study was reviewed and approved by the Medical Ethical Committee Brabant (Tilburg, Netherlands, NL51305.028.15) and all participants provided written informed consent.

### HLA-typing, serological and cellular responses to whole-cell *C. burnetii* at inclusion

HLA typing was performed at the HLA laboratory at the Laboratory of Translational Immunology at the UMC Utrecht, the Netherlands, by Next Generation Sequencing and the resulting HLA-A, HLA-B, and HLA-DRB1 alleles were assigned to supertype families as described previously (7).

IgG and IgM antibody titers for phase I and phase II *C. burnetii* were determined by immunofluorescence assay (Focus Diagnostics) at the Jeroen Bosch Hospital, ’s-Hertogenbosch, the Netherlands.

Cellular responses were determined by whole blood IFNγ release assay (Q-detect™ IGRA), using lithium-heparin anti-coagulated blood stimulated with *C. burnetii* antigen (heat killed Cb02629, Wageningen Bioveterinay Research, lot 14VRIM014) and appropriate positive and negative controls, as described previously (7). In addition to IFNγ ELISA, multiplex cytokine analysis of whole blood stimulation supernatants for IFNγ, IL-2, IL-10, TNFα and IL-1β was conducted using the Human Proinflammatory Panel 1 V-Plex assay (Mesoscale Discovery), according to the manufacturers’ recommendations. Of note, the V-Plex assay uses a different standard than the Q-detect ELISA, resulting in an approximately 20-fold difference in calculated IFNγ concentrations.

### Analysis of *C. burnetii* epitope-specific T-cell responses

Antigen-specific T-cell responses to *C. burnetii* were determined by enzyme-linked immune spot (ELISpot) assay using previously published 50 broadly promiscuous HLA class II epitope clusters (Supplementary Table S1) and 65 HLA class I epitopes (Supplementary Table 2). As described previously (7), these 115 epitopes were derived by immunoinformatic prediction using the iVAX toolkit developed by EpiVax (http://epivax.com/immunogenicity-screening/ivax-web-based-vaccine-design) (25, 26) from two sets of *C. burnetii* antigens: type IV secretion system substrates (T4SS) expected to elicit CD8 responses and known sero-reactive *C. burnetii* antigens based on antibody responses in humans and mice.

Two different ELISpot assays were employed to facilitate detection of central memory T-cell responses (cultured ELISpot) and effector memory T-cell responses (standard or direct ELISpot) (13). ELISpot was conducted using freshly isolated peripheral blood mononuclear cells (PBMCs) from lithium-heparin anti-coagulated blood, using Leukosep tubes prefilled with Ficoll (Greiner BioOne) according to the manufacturer’s recommendations. ELISpot assays were conducted based on a published protocol, using MultiScreen IP filter plates (Merck Millipore) and a human IFN-γ ELISPOT antibody and reagent set (Diaclone) to detect responses to individual peptides in quadruplicate (final concentration 2 μg/ml per peptide, 0.02% DMSO) (7). Plates were scanned on an AID Classic reader system and spot forming units counted using the AID ELISpot software v7.0 (both AID Diagnostika GmbH). Statistical analyses were carried out using GraphPad Prism software (version 7).

For detection of *C. burnetii*-specific central memory T-cell responses and to increase sensitivity for low frequency antigen-specific T-cells, ELISpot was preceded by antigen-specific T-cell expansion with peptide pools (7). Based on cell availability, a median of 41,000 cells per expansion culture (interquartile range (IQR) 32,000-51,000) were plated per replicate well for chronic Q fever subjects and data were analyzed as described previously using three combined threshold criteria (7): Cultured ELISpot peptide re-stimulation responses were defined as positive when they were (i) significantly higher than spot counts in matched negative control wells from the same expansion culture by one-way ANOVA with Holm-Šídák multiple comparison correction post-hoc test, reached (ii) a stimulation index of at least 2 above the matched negative control wells and (iii) an absolute cut-off of >10 SFU/well.

For direct ELISpot, epitope-specific HLA class II responses were evaluated *ex vivo* without prior culture or expansion for all chronic individuals with a sufficiently large number of PBMCs available (n=13), and for a subset of convalescent individuals (n=11). Due to the expected lower pre-curser frequency in fresh PBMCs and based on cell availability, a median of 215,000 cells (interquartile range (IQR) 115,000-331,000) were plated per replicate well for chronic Q fever and convalescent subjects. In addition to peptide stimulation, duplicate wells of PBMCs were also stimulated with whole cell heat-killed *C. burnetii* antigen (strain Cb02629) at the same concentration as used for whole blood stimulations in Q-detect™. The threshold criteria for a positive response in the direct ELISpot assay were (i) spot counts significantly higher than those in matched negative control wells from the same donor by one-way ANOVA with Holm-Šídák multiple comparison correction post-hoc test, (ii) responses that reached a stimulation index of at least 2 above the matched negative control wells and (iii) an absolute cut-off of 10 SFU/million cells.

## Acknowledgements

We would like to thank all chronic Q fever patients as well as the volunteers from the village of Herpen, The Netherlands, for their participation in this study. We acknowledge the patient organizations Q-support and Q-uestion and the following physicians for their assistance in recruiting chronic Q fever patients into this study: M van Kasteren (Elisabeth-TweeSteden Hospital Tilburg) and A. Olde Loohuis. P. Hindocha is acknowledged for assistance with HLA supertype assignment.

This research was supported by contract HDTRA1-15-C-0020 from the US Defense Threat Reduction Agency (www.dtra.mil), awarded to Massachusetts General Hospital (MGH; MCP Lead Principal Investigator); work by authors at other institutions was supported by subcontracts under the prime contract award to MGH. The funder had no role in study design, data collection, analysis or interpretation of the data, the preparation of the manuscript, or the decision to submit the work for publication.

AG is a senior officer and shareholder and AS is an employee of Innatoss Laboratories B.V., which provides diagnostic screening for Q fever. ADG and WM are senior officers and shareholders, and LM and GR are employees of EpiVax, Inc., a company specializing in immunoinformatic analysis. Innatoss Laboratories B.V. and EpiVax, Inc., own patents to technologies utilized by associated authors in the research reported here. The remaining authors declare that the research was conducted in the absence of any commercial or financial relationships that could be construed as a potential conflict of interest.

AS, PMR, MCP, AES and AG formulated research goals; AS and AG designed experiments; EH collected clinical data; CPBR supported patient recruitment; SRP advised on patient selection and reviewed clinical data; AS conducted experiments and analyzed data; AG, WDM and ADG contributed vital reagents and computing tools; LM and GR performed immunoinformatic epitope predictions and selection; GR analyzed HLA supertypes; AS and AG interpreted the data and wrote the manuscript; GR, LM, EH, CPBR, PMR, SRP, WDM, ADG, MCP and AES discussed data and critically revised the manuscript. All authors read and approved the final manuscript.

